# Diet-induced modifications to human microbiome reshape colonic homeostasis in irritable bowel syndrome

**DOI:** 10.1101/2021.08.15.456374

**Authors:** Ayelet Pearl, Hadar Bootz, Ehud Melzer, Efrat Sharon, Shlomi Abuchatzera, Sivan Amidror, Elana Aretz, Irit Shoval, Orly Yaron, Stephen Malnick, Nissan Yissachar

**Affiliations:** The Goodman Faculty of Life Sciences, Bar-Ilan University, Ramat-Gan, 5290002, Israel; Bar-Ilan Institute of Nanotechnology and Advanced Materials, Bar-Ilan University, Ramat-Gan, 5290002, Israel; Department of Gastroenterology and Liver Disease, Kaplan Medical Center; Department of Internal Medicine, Kaplan Medical Center; Hadassah Medical Center, the Hebrew University, Jerusalem, Israel

## Abstract

Changes in microbiome composition have been associated with a wide array of human diseases, turning the human microbiota into an attractive target for therapeutic intervention. Yet clinical translation of these findings requires the establishment of causative connections between specific microbial taxa and their functional impact on host tissues. Here, we colonized gut organ cultures with longitudinal microbiota samples collected from newly-diagnosed and therapy-naïve irritable bowel syndrome (IBS) patients under low-FODMAP (fermentable Oligo-, Di-, Mono- saccharides and Polyols) diet. We show that post-diet microbiota regulates intestinal expression of inflammatory and neuro-muscular gene-sets. Specifically, we identify *Bifidobacterium adolescentis* as a diet-sensitive pathobiont that alters tight junction integrity and disrupts gut barrier functions. Collectively, we present a unique pathway discovery approach for mechanistic dissection and identification of functional diet-host-microbiota modules. Our data support the hypothesis that the gut microbiota mediates the beneficial effects of low-FODMAP diet, and reinforce the potential feasibility of microbiome based-therapies in IBS.

## Introduction

Irritable Bowel Syndrome (IBS) is a functional disorder of the gastrointestinal (GI) tract, which causes chronic abdominal pain, changes in bowel habits and frequent bloating, resulting in a significant impairment in the quality of life (Enck et al., 2016). Several IBS subtypes have been classified according to the Rome IV criteria, including diarrhea- or constipation-predominant IBS (IBS-D and IBS-C, respectively), IBS with mixed bowel habits and unclassified IBS (Palsson et al., 2016). Several anatomical systems have been associated with IBS pathogenesis, including the intestinal muscle layers, the enteric (ENS) and the central (CNS) nervous systems and the intestinal immune system. However, the etiology of IBS remains poorly understood, partially due to its pathophysiological heterogeneity and the absence of robust animal models. Significantly, changes in gut microbiota composition have emerged as modulators of disease pathophysiology (Mars et al., 2020), and dietary habits, which influence gut microbiota composition, have shown significant effects on disease manifestations (Moayyedi et al., 2020). The most effective treatment for IBS is a 6-week low FODMAP (fermentable Oligo-, Di-, Mono-saccharides and Polyols) diet (Altobelli et al., 2017; Staudacher and Whelan, 2017; Staudacher et al., 2014). Although a low-FODMAP diet was shown to alter gut microbiome composition (Halmos et al., 2015; Hustoft et al., 2017), no clear microbiota signature has emerged for diet-responding patients (Bennet et al., 2018; McIntosh et al., 2017).

Determining whether diet-induced alterations to microbiome composition have functional consequences on the local gut tissue remains a major challenge. Such studies cannot be faithfully accomplished in tissue culture monolayers, or even in 3D organoids that lack the anatomical complexity and heterogeneity of the intact intestine. On the other hand, *in vivo* gnotobiotic models are generally limited for throughput due to technical constraints related to expense, time and labor-intensiveness. To overcome these challenges, we recently developed a gut organ culture system (Yissachar et al., 2017). This system maintains the naive intestinal tissue structure and cellular complexity, yet, as in simple *in vitro* assays, provides tight experimental control over the luminal content, and is ideal for analyzing the effects of rapid luminal perturbations on gut-residing cells. In the current study, we leveraged the gut organ culture system to investigate colonic responses to whole microbiota communities collected from IBS patients, pre- and post-low-FODMAP diet. We show that post-diet microbiota rapidly alters colonic gene expression, and describe novel associations between dietary-altered microbial taxa and differentially expressed intestinal genes. Intriguingly, our findings mechanistically link low-FODMAP diet and its associated microbiome with the regulation of intestinal barrier functions in IBS.

## Results

### A longitudinal cohort of diarrhea-predominant IBS throughout low-FODMAP diet

To examine the effects of low-FODMAP diet on gut microbiome composition and colonic homeostasis, we established a longitudinal cohort of newly-diagnosed and therapy-naïve IBS-D patients (n=10; fulfilled the Rome IV criteria; 6-weeks follow-up per patient), with longitudinal clinical evaluations and microbiome sampling throughout 6-weeks of low-FODMAP diet (Fig. 1A). Full medical history was obtained for all patients. Anthropometrics, clinical and subjective evaluations of disease status (IBS-SSS), dietary consultation, and fecal sample collection and evaluation (Bristol stool score) were performed at time 0 and at 3 and 6 weeks under low-FODMAP diet (Fig. 1A). Clinical responses were consistent with previous reports (Gearry et al., 2016), with 70% of patients (n=7) positively responding to the diet (overall improvement in disease manifestations, particularly reduced abdominal pain duration and increased bowel habits satisfaction) (Fig. S1A). Similarly, significant improvement in Bristol stool score was observed for responding patients, but not for non-responders (Fig. S1B). Interestingly, we noticed a significant weight loss in all patients with mean change of -1.74kg (SE=±0.27, or 2.5% reduction from initial weight, screening to end-phase; weight loss was independent of clinical response) (Fig. 1B). Overall, the clinical characteristics of this longitudinal cohort are comparable with previously described responses to low-FODMAP diet in IBS, and as such, is a suitable representative cohort to study the interplay among diet, gut microbiome and colonic responses.

**Fig. 1.**
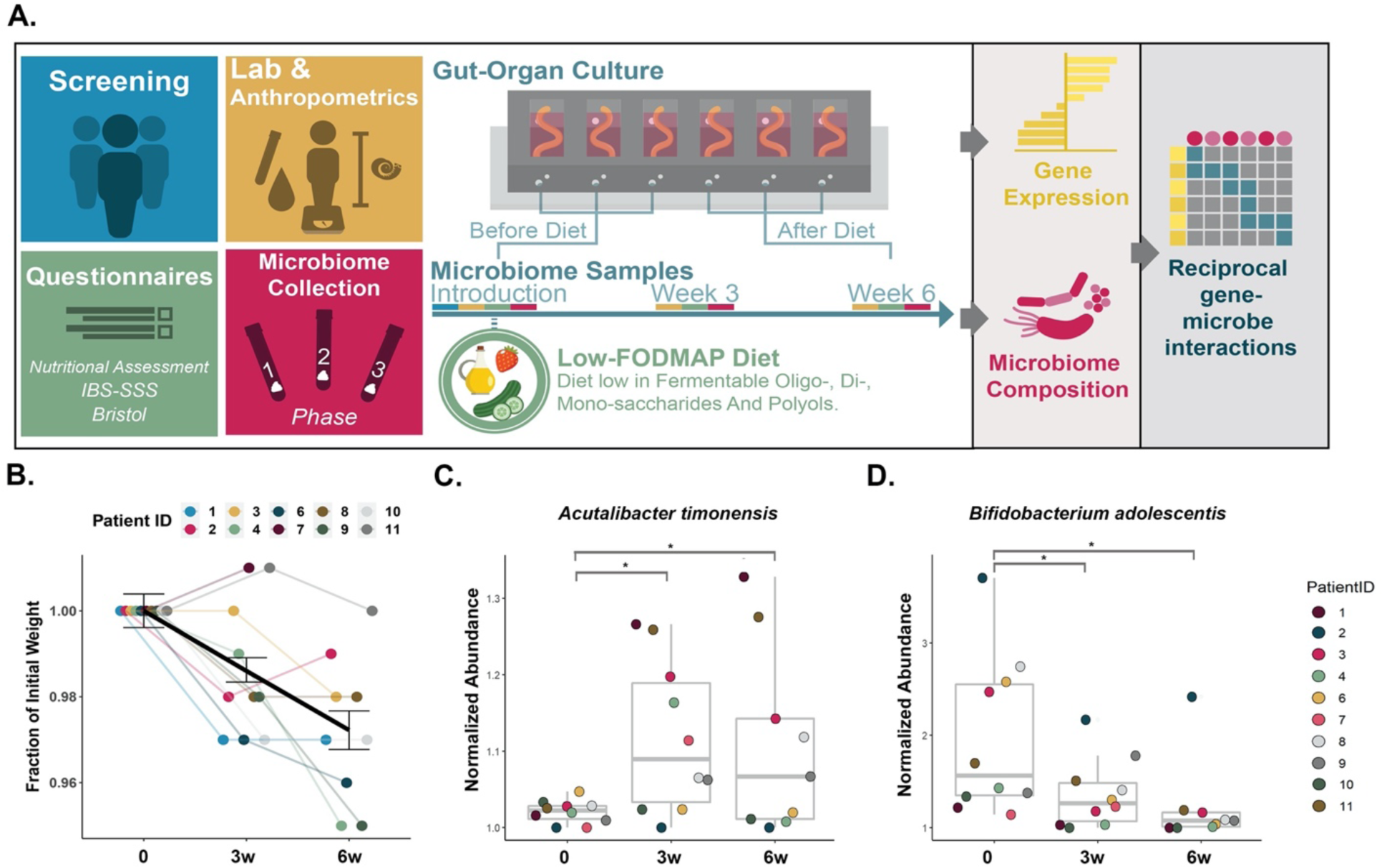
Low-FODMAP diet modifies gut microbiome composition in IBS-D patients. **(A)** Experimental outline: a longitudinal cohort of newly-diagnosed and therapy-naïve IBS-D patients included clinical evaluations and microbiome sampling throughout 6-weeks of low-FODMAP diet. Microbiome composition was determined by 16s rRNA gene sequencing. Functional impact of post-diet microbiota was determined by *ex vivo* colonization of colon organ cultures with patient’s microbiota samples, pre- and post-diet. Colonic gene expression was determined using bulk RNA sequencing, followed by predictions of reciprocal associations between specific microbial taxa and differentially expressed host genes. **(B)** Weight loss was observed in IBS patients following low-FODMAP diet. Weight is represented as fraction of the initial (pre-diet) weight, at mid-phase (3w) and end-phase (6w) of the diet. Colored lines represent individual patients, legend shows patient ID by color); black line represents average weight change with standard error bars. Statistical significance was determined by Student’s T-test, \**p*<0.05; \*\**p*<0.001. **(C-D)** Normalized abundance of *A.timonensis* **(C)** and *B.adolescentis* **(D)** in patients’ gut microbiota at initial (0w), mid-phase (3w) and end-phase (6w) of low-FODMAP diet. Legend: patient ID by color. Statistical significance was determined by Wald test, \*\**p*<0.01.

### Low-FODMAP diet modifies gut microbiome composition

Gut microbiome composition was analyzed by 16S rRNA gene sequencing of patients’ fecal samples collected at 0, 3 and 6 weeks under low-FODMAP diet. In agreement with previous studies (Bennet et al., 2018; Halmos et al., 2015; McIntosh et al., 2017; Moayyedi et al., 2020; Pittayanon et al., 2019; Staudacher et al., 2014), we did not detect statistically significant changes in per-sample bacterial richness (alpha diversity) or between-sample (beta) diversity, throughout the diet course. Furthermore, analysis of Euclidean distances between samples indicated that samples were clustered by patient, then by treatment, suggesting that the effects of low-FODMAP diet on microbiome composition are secondary to inter-individual diversity (Fig. S1C). Yet, analysis of differentially abundant bacterial taxa revealed that the post-diet microbiome is characterized by increased levels of *Acutalibacter timonesis* and *Oscillibacter sp900066435* species (Fig. 1C and Fig. S2C), and conversely, decreased abundance of several bacterial species, most notably *Eubacterium ventriosum*, *Clostridium disporicum* and *Bifidobacterium adolescentis* (Fig. 1D and Fig. S2C). Among others, genus-level comparison detected reduced abundance of *Bifidobacterium* and *Eubacterium* following low-FODMAP diet (Fig. S2B), in agreement with previous reports (Bennet et al., 2018; Staudacher et al., 2012, 2017). Thus, compared with pre-diet microbiome, low-FODMAP diet significantly altered the relative abundance of specific microbial taxa in IBS-D patients.

### Post-diet microbiota modifies colonic gene expression *ex vivo*

To determine whether diet-induced alterations to microbiome composition functionally affect colonic gene expression, intact colon tissues were dissected from specific-pathogen free (SPF) mice and connected to the gut organ culture system, as previously described (Duscha et al., 2020; Yissachar et al., 2017). Gut organ cultures were colonized *ex vivo* with fecal samples collected from the first 4 patients that successfully completed the clinical phase (3 responders and 1 non-responder), pre- and 6 weeks post-diet, at equivalent bacterial loads (10^7^ CFUs/mL) (n=4 patients, pre- and post-diet, in duplicates, for a total of 16 samples). To characterize immediate-early gene expression, bulk RNA sequencing was performed on whole-tissue colon samples at 2h post-colonization. Differential gene expression analysis, comparing colonic transcriptional responses to post-diet versus pre-diet microbiota, revealed rapid increased expression of 108 transcripts and decreased expression of 98 genes (fold change>2 and *p*<0.05; Fig. 2A; Table S1). Interestingly, Gene Ontology (GO) analysis indicated that among the genes induced by post-diet microbiota were those implicated in enteric neuronal and muscle functions, including tachykinin receptors *TacR1* and *TacR2*, cholecystokinin receptor A *Cckar*, neuropeptide Y receptor *Npy4r*, neuromedin U receptor *NmuR1*, alpha-adrenergic receptor *Adra1b*, and type C natriuretic peptide *Nppc*. We also identified extracellular signaling molecules *Ccn3* and *Ccn6* (which mediate cell-matrix interaction during angiogenesis and wound healing), and the resistin-like molecule-α *Retnla* (which encodes the antimicrobial protein RELM-α) (Fig. 2A). Among the genes suppressed by post-diet microbiota, we identified genes expressed by various immunological cellular subsets, including the pro-inflammatory cytokine subunit *Il12b* (macrophages and dendritic cells), the B cell chemoattractant *Cxcl13* (macrophages), the B cell antigen receptor component *CD79a*, the complement receptor 2 gene *Cr2* (B cells), the neutrophils/granulocytes marker *Ly6g*, the TNF-receptor superfamily member *Tnfrsf8* (expressed on activated T- and B-cells), and the disintegrin and metalloproteinase family member *Adamts15*, highly expressed on intestinal type-3 innate lymphoid cells (ILC3) (Fig. 2A). These results imply that post-diet microbiota exerts rapid immunosuppressive effects on a wide array of colonic immunocyte lineages.

**Fig. 2.**
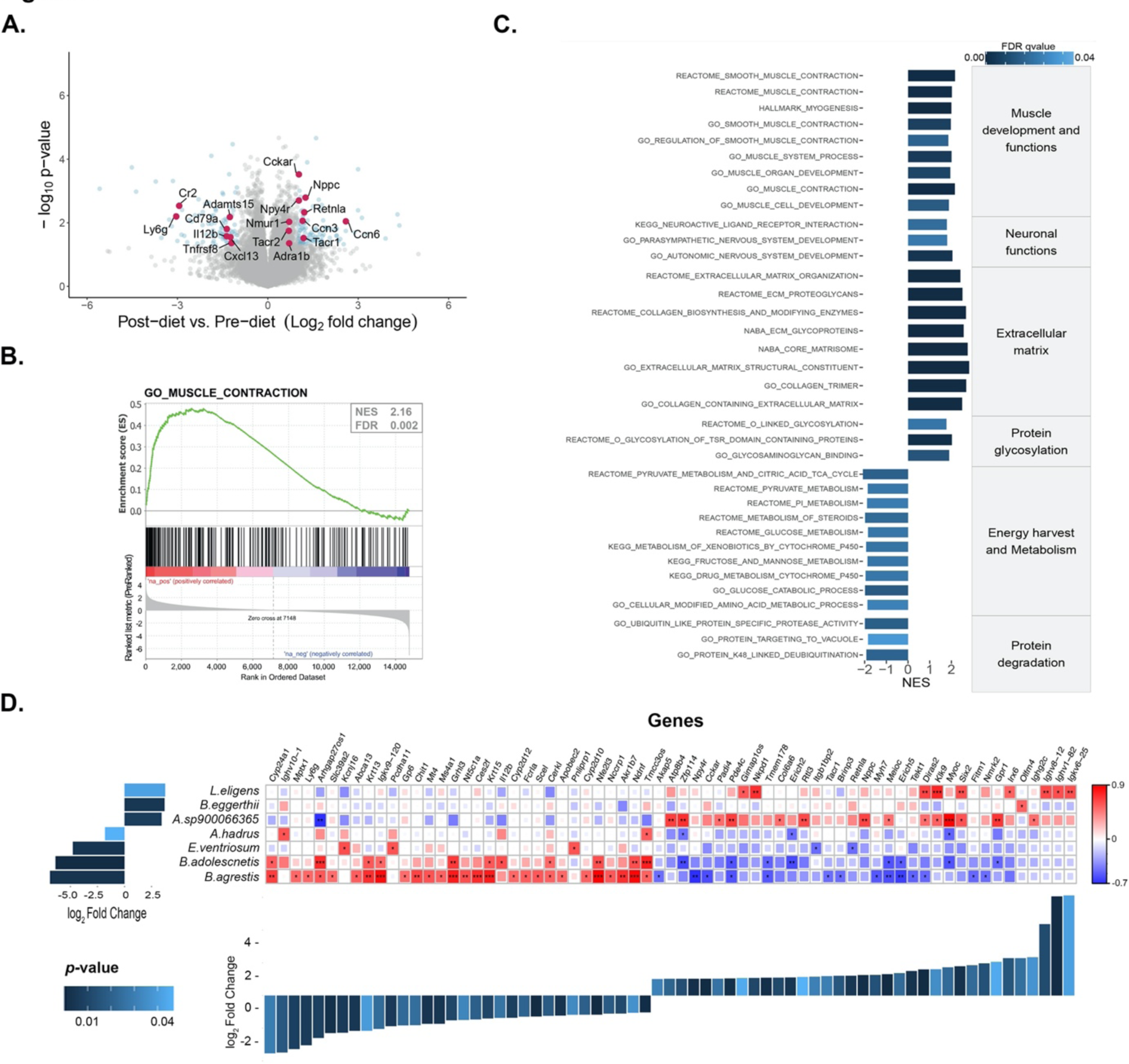
Post-diet microbiota modifies colonic gene expression *ex vivo*. **(A)** Volcano plot indicating changes in gene expression comparing colon organ cultures infused for 2h with post- diet (6w) vs. pre-diet microbiota (n=4 for each condition, in duplicates, total of 16 samples were sequenced). Transcripts significantly up- or downregulated (blue; fold change > 2, *p*<0.05), as well as selected genes of interest (red), are highlighted. **(B-C)** Gene-set enrichment analysis (GSEA) identified gene pathways significantly up- or downregulated following gut colonization with post- vs. pre-diet microbiota. **(B)** Enrichment score plot for a representative gene ontology pathway, and **(C)** Top significant pathways selected from the hallmark, Curated Canonical Pathways and GO gene sets databases, grouped into categories. The normalized enrichment scores (NES) and FDR *q*-values (<0.05; scale bar) are indicated. **(D)** Associations (pairwise Pearson correlation) between differentially expressed colonic genes (fold change>2, *p*<0.05) in colons infused with post- vs. pre-diet microbiota and the relative abundance of microbial species (fold change>2, *p*<0.05) in the corresponding microbiota samples. Positive and negative Pearson’s correlation scores are represented by red to blue scale bar (right). Significant correlations are indicated by * (*p*<0.05), ** (*p*<0.01), *** (*p*<0.001). Bar plots represent changes in bacterial abundance (left) and colonic gene expression (bottom), post- vs. pre-diet. Scale bar (left) indicates *p* values for both bar plots.

Intriguingly, an unbiased gene set enrichment analysis (GSEA) revealed that post-diet microbiota significantly induced transcriptional pathways related to muscle contraction, development and function (Fig. 2B-C), as well as pathways related to neuronal activity, modifications of the extracellular matrix, and protein glycosylation (Fig. 2C). As the gastrointestinal muscle and the enteric nervous system are highly involved in IBS pathophysiology, these initial findings may suggest a role for post-diet gut microbiota in directing neuro-muscular functions in IBS. In contrast, gene programs that control energy harvest and metabolism (including mitochondrial pyruvate metabolism and citric acid cycle), and several protein degradation pathways, were down-regulated compared with tissues colonized with pre-diet microbiota (Fig. 2C).

### Significant associations between specific microbial taxa and colonic gene expression

We next asked whether microbiota-induced alterations in colonic gene expression could be associated with specific bacterial taxa, based on the known composition of the microbiota introduced into each colon organ culture. Pearson-correlation analysis was used to connect pre-and post-diet microbiome composition from each of the 4 patients (infused into the gut cultures), with the corresponding colonic transcriptional profiles induced following *ex vivo* colonization. Out of 160 unique bacterial taxa identified in these samples, 7 showed a statistically significant correlation (*p*<0.05) with 67 host genes (35 genes positively correlated, and 32 genes negatively correlated) (Fig. 2D). For example, luminal abundance of the enterobacterium *Buttiauxella agrestis* showed a significant positive correlation with colonic expression of many immunocyte-related genes (including *Fcrla* (B cells), *Ly6g* and *Abca13* (neutrophils/ granulocytes), *Chit1* and *Akr1b7* (macrophages) and *Scel* (ILC2)) and a negative correlation with several neuronal and muscle-related genes, including the neuropeptides receptors *Npy4r*, *Cckar* and *Tacr1* (Fig. 2D). In addition, the abundance of the T helper-17 (Th17)-inducing microbe *B. adolescentis* (Tan et al., 2016) was positively correlated with colonic expression of several epithelial-related genes (i.e. *Krt13* and *Krt15*) and immunological genes (i.e. *Il12b*) (Fig. 2D). Interestingly, *Il12b* encodes the common p40 subunit of the pro-inflammatory cytokines IL-12 and IL-23. *Il12b* is expressed by dendritic cells and macrophages and promotes the pro-inflammatory activities of Th1/Th17 cells. Overall, these findings predict a functional interplay between particular dietary-modulated bacterial taxa and specific intestinal genes.

### *B. adolescentis* disrupts epithelial tight junction integrity and gut barrier functions

Our analysis revealed an association between luminal *B. adolescentis* abundance and the expression of barrier-related genes (Fig. 2D), including keratins *Krt13* and *Krt15* (which encode intermediate filaments that maintain epithelial structural integrity), *Il12b* (which is rapidly induced following disruptions to the gut epithelial barrier (Eftychi et al., 2019)), as well as *Ndnf* and *Myoc* (both of which are associated with regulation of cell adhesion, per GO:0045785 pathway). Like other Th17-inducing microbes, *B. adolescentis* possesses the ability to tightly associate with the intestinal epithelium (Atarashi et al., 2015; Tan et al., 2016). Moreover, low- or high-FODMAP diets were shown to ameliorate or exacerbate, respectively, intestinal inflammation and barrier dysfunction in rats (Zhou et al., 2018). In our study, low-FODMAP diet significantly reduced *B. adolescentis* abundance, compared to its pre-diet levels (Fig. 1D). Based on these lines of evidence, we hypothesized that in the context of IBS, *B. adolescentis* may disrupt epithelial cell integrity and gut barrier functions.

To test this hypothesis, we co-cultured colonic epithelial cells (CaCo-2) with *B. adolescentis*, or with an equivalent load of *B. fragilis* (a non-Th17-inducing symbiont (Mazmanian et al., 2008)). *B. adolescentis*, but not *B. fragilis*, induced a rapid (i.e. within 4h) and significant suppression of *Tjp1* expression (p<0.01), which encodes the tight junctions (TJs) adaptor protein ZO-1 (Fig. 3A). In addition, *B. adolescentis* significantly induced the expression of *Cldn2* (p<0.001), which encodes a pore-forming component of the Claudin family of TJs proteins (Tsai et al., 2017) (Fig. 3A). Immunofluorescence staining of CaCo-2 cells identified marked decrease (p<0.0001) in ZO-1 protein expression levels at the epithelial TJs in response to *B. adolescentis* (Fig. 3B-C). In agreement, dynamic measurements of trans-epithelial electrical resistance (TEER) for CaCo2 monolayers revealed that the introduction of *B. adolescentis*, but not *B. fragilis*, impaired epithelial barrier integrity (p<0.0001), as indicated by rapid reduction in TEER values (Fig. 3D-E). To exclude possible laboratory-dependent variations in bacterial stocks, we repeated the TEER experiments with *B. adolescentis* obtained from two different sources (laboratory collection per (Tan et al., 2016)) and DSMZ) and obtained identical results (Fig S3A-B). Interestingly, addition of material secreted by *B. adolescentis* had no impact on epithelial TEER (Fig. 3E), suggesting that the regulation of barrier function by *B. adolescentis* may be dependent on bacterial adhesion to the gut epithelium. In contrast, addition of heat-killed *B. adolescentis* significantly impaired TEER barrier index compared with untreated cells, but to a lesser extent than the live microbe, suggesting that bacterial components mediate epithelial barrier disruption in a contact-dependent manner (Fig. 3F).

**Fig. 3.**
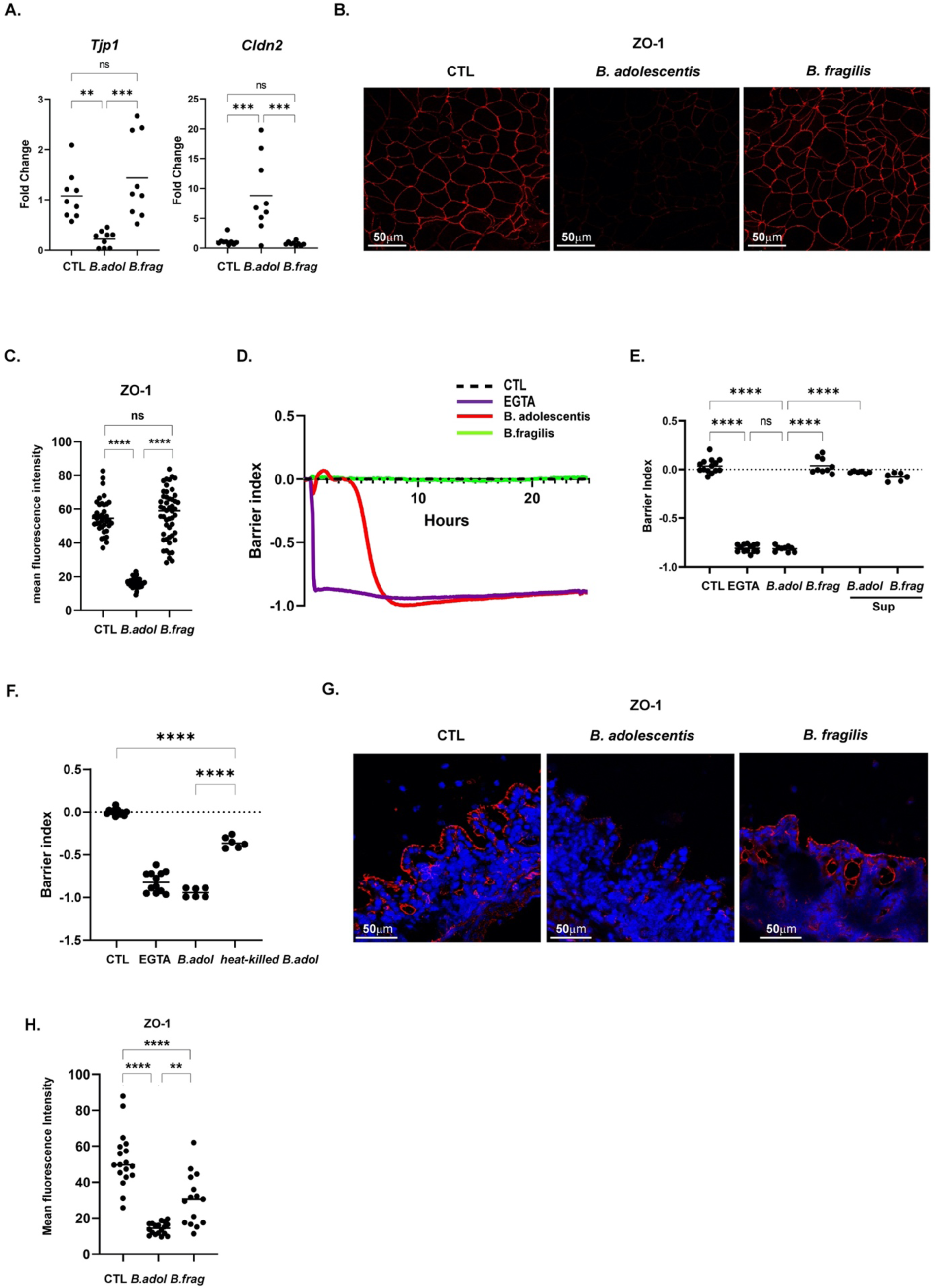
*B.adolescentis* disrupts TJ integrity and epithelial barrier functions. **(A)** Gene expression assessed by RT-PCR in CaCo-2 cells co-cultured for 4h with *B.adolescentis* or *B.fragilis*. CT values are normalized to the reference gene, EEF2. Statistical significance was determined by ANOVA, *** p< 0.001; ** *p* < 0.01; ns - not significant. Data acquired by three independent experiments. **(B-C)** Confocal microscopy images stained for ZO-1 **(B)** and quantification of Mean Fluorescence Intensity (MFI) **(C)** of CaCo-2 cells following different treatments. **(D-F)** Normalized TEER values vs. time **(D)** and after 24h **(E-F)**. **(G-H)** Confocal microscopy images of ZO-1 (red, nuclei were counterstained with DAPI, blue) **(G)** and quantification of Mean fluorescence intensity (MFI) **(H)** in colon organ cultures at 4h post-colonization with *B.adolescentis* (B.adol), *B.fragilis* (B.frag) or sterile medium (CTL), using the gut organ culture system.

To further examine whether *B. adolescentis* disrupts the epithelial barrier of intact colon tissues, we infused equivalent loads of *B. adolescentis* or *B. fragilis* into the lumen of colon organ cultures for 4h. Here again, *B. adolescentis* compromised TJ integrity, as indicated by a significant decrease (p<0.0001) in epithelial ZO-1 protein staining in colon organ cultures (Fig. 3G-H).

These results prompted us to examine whether *B. adolescentis* disrupts epithelial barrier integrity and increases gut permeability *in vivo*. Littermates from C57BL/6 mice reared in a SPF facility were pre-treated with broad range antibiotics (vancomycin, metronidazole, neomycin, and ampicillin) in the drinking water for 1 week, followed by daily oral gavage with purified cultures of *B. adolescentis*, *B. fragilis*, or PBS (as a control) for 3d. A significant increase in the uptake of orally fed fluorescein isothiocyanate (FITC)–dextran into the systemic circulation was observed in mice following oral gavage with *B. adolescentis*, but not *B. fragilis*, compared to PBS-fed littermates controls, indicating impairment of gut barrier (p=0.013) and increased gut permeability (Fig. 4A).

**Fig. 4.**
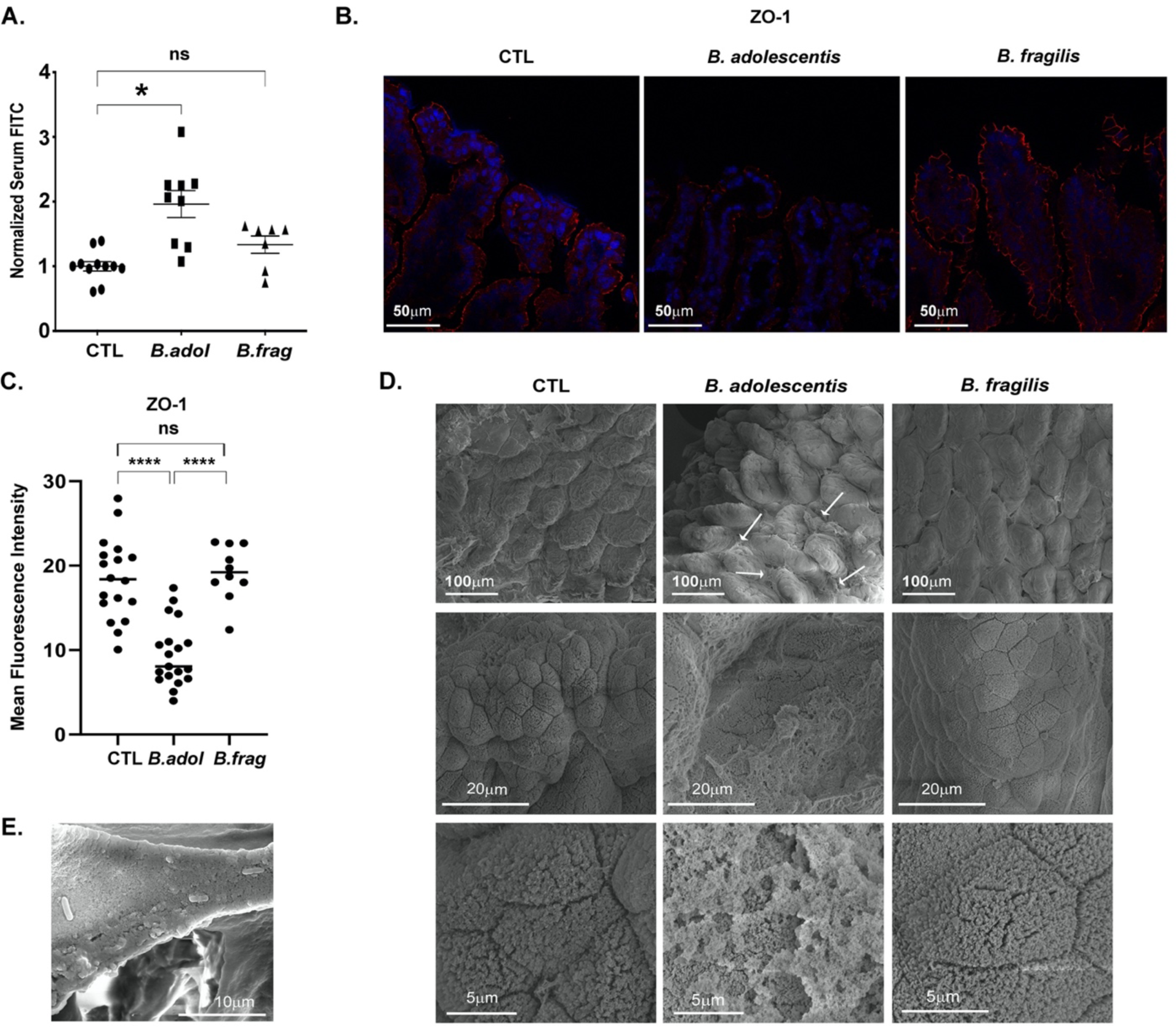
*B.adolescentis* disrupts TJ integrity and increases gut permeability *in vivo*. **(A)** Serum levels of orally administrated FITC-dextran from mice following oral gavage of *B.adolescentis* (B.adol), *B.fragilis* (B.frag) or sterile PBS (CTL). **(B-C)** Confocal microscopy images of ZO-1 (red, nuclei counterstained with DAPI, blue) **(B)** and quantification of Mean Fluorescence Intensity (MFI) **(C)** in small intestine of mice following oral gavage of *B.adolescentis* (B.adol), *B.fragilis* (B.frag) or sterile PBS (CTL). **(D-E)** Representative SEM images of the terminal ileum of mice colonized as indicated (D) and EPS-associated bacterial cells (E). Arrows indicate sites of matrix suspected to be bacterial derived biofilm.

In agreement with the results obtained *in vitro* (Fig. 3B-C) and *ex vivo* (Fig. 3G-H), *in vivo* supplementation of *B. adolescentis* resulted in a substantial disruption to epithelial tight-junction integrity, as indicated by reduced (p<0.0001) immunofluorescence staining of epithelial ZO-1 protein in small intestinal (terminal ileum) tissues (Fig. 4B-C). Interestingly, ZO-1 protein expression in colon tissues was not significantly altered (Fig. S3C-D). We speculate that the dense colonic mucus layer disrupts the association of *B. adolescentis* with the colonic epithelium, thus prevents barrier disruption (Johansson et al., 2008).

We next investigated whether *B. adolescentis* supplementation disturbs the ultrastructure architecture of the intestinal epithelial barrier, using scanning electron microscopy (SEM). In mice orally gavaged with *B. fragilis* or with sterile PBS, the intestinal epithelial layer was intact, clearly defined microvilli were easily observed at the epithelial brush border, and epithelial adherent bacteria were rarely detected (Fig. 4D). In contrast, following oral gavage of *B. adolescentis,* we detected large quantities of a substance that mostly resembles bacterial-derived exopolysaccharide (EPS), recently shown to be produced by *B. adolescentis* and other members of the *Bifidobacterium* genus (Fanning et al., 2012; Hughes et al., 2017; Yu et al., 2019). This substance accumulated on and between the small intestinal villi, covering and disrupting the gentle structure of the epithelial brush border (Fig. 4D), and associated bacteria were frequently found (Fig. 4E).

Collectively, these results indicate that *B. adolescentis* disrupts TJ integrity and epithelial barrier functions in epithelial cell cultures, in intestinal organ cultures, and in mice.

## Discussion

Diet is one of the most potent factors that determine the gut microbiome structure, and emerging dietary interventions are now designed to reshape the gut microbiome into a healthier configuration (Rothschild et al., 2018; Zeevi et al., 2015). Although low-FODMAP diet is a relatively effective treatment for IBS, it benefits only some patients, and cannot be used as a long-term treatment due to inherent nutritional deficiencies and low compliance. Here, we provide evidences that low-FODMAP diet alters gut microbiome composition in newly-diagnosed and therapy-naïve IBS patients, and that post-diet microbiota modifies colonic gene expression in gut organ cultures. Furthermore, we show that low-FODMAP diet diminishes luminal abundance of *B. adolescentis*, which we identify as a potent disruptor of epithelial TJ integrity and intestinal barrier functions. Our data support the hypothesis that the gut microbiota mediates the beneficial effects of low-FODMAP diet, and reinforce the potential feasibility of microbiome based-therapies in IBS (i.e., using fecal microbiota transplants (El-Salhy et al., 2020) or targeted/rational modifications to the microbiome).

The importance of longitudinal sampling and analysis was recently demonstrated for IBS subtypes compared with healthy individuals, revealing a role for the gut microbiota and its related metabolites in regulating IBS pathogenesis (Mars et al., 2020). The effect of low-FODMAP diet on gut microbiome composition in IBS patients was analyzed in several recent studies (Bennet et al., 2018; Halmos et al., 2015; McIntosh et al., 2017; Moayyedi et al., 2020; Pittayanon et al., 2019; Staudacher et al., 2014), revealing differentially abundant bacterial taxa (i.e. *Actinobacteria*, *Bifidobacterium* and *Eubacterium*), but without substantial alterations to overall alpha- and beta-diversity. Our longitudinal analysis complements and extends these findings with the identification of specific bacterial species affected by low-FODMAP diet, in particular, *Bifidobacterium adolescentis* among the bacterial taxa whose abundance was most prominently decreased following diet. Interestingly, reduced abundance of *B. adolescentis* was recently observed in quiescent inflammatory bowel diseases (IBD) patients under low-FODMAP diet (Cox et al., 2020). In addition, low levels of *B. adolescentis* were linked to greater weight loss in obesity treatment program (Santacruz et al., 2009), and were associated with ‘good’ personalized diet that lowers post-meal blood glucose (Zeevi et al., 2015). As members of the *Bifidobacterium* genus possess a unique ability to ferment carbohydrates via the fructose-6-phosphate phosphoketolase (F6PPK) pathway, the decline in *B. adolescentis* levels may result from reduced oligosaccharide availability under a low-FODMAP diet (as well as in other beneficial dietary interventions).

Our work took these findings a significant step further by assessing the impact of the transformed post-diet microbiota on colonic gene expression and function. Despite substantial advances in analyzing microbiome composition and metabolic activity, the ability to determine whether changes to microbiome composition functionally impact intestinal homeostasis remains a major challenge. Here, we addressed this challenge by colonizing gut organ cultures with patient-derived microbiota, pre- and post-diet. With this system, we could control the input (i.e. luminal content) as in simple cell culture systems, with the advantage of determining functional changes in an *ex vivo* model of the gut that more closely mimics the *in vivo* organ. This enables determination of direct correlations between specific bacterial taxa and colonic changes, which are not possible in complex *in vivo* models. For example, the association of *Buttiauxella agrestis* abundance with colonic expression of immunological, neuronal and muscle-related genes may potentially link the high abundance of *Enterobacteriaceae* previously observed in IBS patients (Pittayanon et al., 2019) with immune activation and alterations to neuro-muscular systems. In addition, we predicted, and experimentally validated, a functional interplay between luminal *B. adolescentis* abundance and enteric expression of epithelial and immunological genes, collectively suggesting that dietary-modulated *B. adolescentis* may drive gastrointestinal barrier dysfunctions in IBS.

Disruptions to the gut epithelial barrier (a ‘leaky’ gut) may lead to severe pathological conditions including infection and inflammation. For example, gut barrier dysfunction was linked to autoimmune reactions in systemic lupus erythematosus (SLE) (Manfredo Vieira et al., 2018), hyperglycemia (Thaiss et al., 2018), IBD (Buckley and Turner, 2018; Khor et al., 2011; Luissint et al., 2016), and Celiac disease (Clemente et al., 2003). In IBS, epithelial barrier dysfunction and increased gut permeability are a hallmark of the disease, and are likely to predispose patients to abnormal immune activation, abdominal pain, and other IBS-related symptoms (Aguilera-Lizarraga et al., 2021; Dunlop et al., 2006; Martínez et al., 2012, 2013, 2017; Piche et al., 2009).

A potential role for the gut microbiota in regulating barrier functions in IBS was demonstrated by transplantation of whole microbiota samples from IBS-D patients into germ-free mice, resulting in intestinal barrier dysfunction, as well as immune activation, increased gastrointestinal transit time and anxiety-like behavior (De Palma et al., 2017). Similarly, the impact of low and high FODMAP diets on gut microbiome composition and barrier functions was recently demonstrated in rats, where high-FODMAP diet increased colonic permeability and expression of pro-inflammatory cytokines, and low-FODMAP diet ameliorated these responses (Zhou et al., 2018). Yet, no specific bacterial species were related to these phenotypes. Our discovery that *B. adolescentis* disrupts gut barrier functions (possibly via contact-dependent disruptions to TJ integrity), and that its abundance is diminished under low-FODMAP diet in human IBS patients, may provide a mechanistic explanation to these observations.

Intriguingly, *B. adolescentis* induces the development of pro-inflammatory Th17 cells in the small intestine (Tan et al., 2016). As low-grade inflammation is a major characteristic of the intestinal mucosa of IBS patients (Aguilera-Lizarraga et al., 2019; Barbara et al., 2007; Cenac et al., 2007; Edogawa et al., 2020; Foley et al., 2011; Guilarte et al., 2007; Matricon et al., 2012; Ohman and Simrén, 2010), we propose to further examine a potential connection between post-diet microbiota in general, and *B. adolescentis* in particular, and low-grade intestinal inflammation in IBS patients.

In conclusion, we present a novel experimental framework for investigating how alterations to human gut microbiota impact intestinal tissue function and host-microbiota interactions. We recently demonstrated that the gut organ culture system is a powerful tool for dissecting host responses to specific immunomodulating bacterial strains (Yissachar et al., 2017), to whole microbiota samples collected from human multiple sclerosis patients (Duscha et al., 2020), and to bacterial-derived metabolites (Grosheva et al., 2020). We now show that the combination of the gut organ culture system with clinically relevant human microbiota samples generates a potent pathway discovery platform that enables proceeding from general diet-induced changes in microbiome composition to the identification of specific bacteria that control intestinal gene expression and function. We anticipate that this unique approach will promote future mechanistic discoveries of host-microbiome interactions, and may potentially support the development of personalized microbiome-based therapies for human diseases.

## Materials and methods

### Patients and study design

The study was open labeled and approved by the local ethical committee (LEC, Kaplan Medical Center, Rehovot, Israel). Consecutive patients with diarrhea-predominant-IBS (IBS-D), diagnosed according to Rome IV criteria (Drossman and Hasler, 2016) who were about to start a low-FODMAP diet, were offered to participate in the study. Patients age 18-35 who gave their consent were included if they were diagnosed as having IBS-D for at least 1 year, had an IBS severity scoring system (IBS-SSS) ≥ 175 (FRANCIS et al., 1997), completed a Bristol Stool Scale (Lewis and Heaton, 1997), BMI 18-25 Kgm/M^2^, with normal Hemoglobin, serum aminotransferases, TSH, serum calcium and C-Reactive Protein within 90d prior to day 1. Patients were excluded if they had other significant gastrointestinal diseases, significant systemic disease, including cardiovascular disease, chronic renal failure (GFR<60 ml/min), chronic liver disease or diabetes mellitus, had antibiotic treatment within 3 months of day 1, travelled to locations in South East Asia, South America or Africa within 6 months of day 1, had blood in their stools, weight loss or night sweats or history of food allergy. Screening (up to 14d before day 1) included a full medical history, IBS-directed history and a physical examination. Follow up visits occurred at 3- and 6-weeks following introduction of the Low FODMAP diet. On each of the three visits anthropometric measurements were recorded, including weight (Kg), height (M) & waist circumference (cm) and BMI was calculated. IBSS and Bristol Stool Scale questionnaires were filled and fresh stool was collected and stored at -80^0^ C.

### Mice

C57BL/6J (B6) mice were obtained from Envigo RMS (Israel), and bred in the specific-pathogen-free (SPF) facility at Bar-Ilan University. For gut organ cultures experiments, intestinal tissues were dissected from 14d old littermate mice. All experiments were performed following animal protocols approved by the Bar-Ilan University ethics committee (ethics approval numbers 620918, 110219).

### Bacterial cultures

Bacteria were obtained from DSMZ (Germany) and laboratory stocks were kindly provided by the Mathis/Benoist laboratory at Harvard Medical School (per (Tan et al., 2016)). Anaerobic bacterial strains were incubated overnight in rich liquid medium (2% proteose peptone, 0.5% yeast extract and 0.5% NaCl) supplemented with 250 mg/ml glucose, 250 mg/ml K2HPO4, 50 mg/ml L-cysteine, 5 mg/ml Hemin and 5ul/ml vitamin K1 at 37°C in an anaerobic chamber (90% N2, 5% CO2, 5% H2). For co-culture experiments, bacterial cultures were collected by centrifugation and resuspended in sterile PBS. For evaluation of bacterial load in fecal samples collected from patients, fecal samples were diluted and plated on rich BHI plates, and incubated at 37°C under anaerobic and aerobic conditions. Bacterial load (CFU/gr) was determined by counting microbial colonies following 48h (anaerobic) and 24h (aerobic) culture. For heat-killed bacteria experiments, microbes were incubated at 95°C for 15 min. Live bacteria could not be detected by subsequent anaerobic bacterial cultures.

### Bacterial DNA extraction, amplification and sequencing

DNA was extracted from all fecal samples using the Mobio PowerSoil DNA extraction kit, as described by the manufacturer (MoBio, Carlsbad, CA). DNA concentrations were measured by nanodrop, and 20 ng of DNA was used for 16S library preparation. Library preparation and sequencing was performed by HyLabs Ltd (Israel), as following: 16S libraries for Illumina sequencing were prepared using a two-step PCR protocol. In the first PCR, primers containing tails were used to amplify the V4 region of the 16S rRNA gene. The Access Array primers for Illumina (Fluidigm) are used in the second PCR to add the adaptor and index sequences to each sample. After the second PCR, the reactions were cleaned using Kapa Pure beads (Kappa) to remove unincorporated primers and any primer-dimer produced. The concentration of each library was measured by Qubit (Life Technologies) using the Denovix ds DNA HS assay (Deonvix) and then the samples were combined in equal amounts to a single pool. The pooled sample was run on the Tapestation (Agilent) using a DNA 1000 screentape to determine the size of the pooled libraries. The pooled library was then loaded on the Illumina Miseq and sequenced using a Miseq V2 Kit (500 cycles) to generate 2x250 PE reads. The data were de-multiplexed using Basespace, the Illumina cloud software to generate 2 FASTQ files per sample.

### Analysis of microbiome composition

FASTQ files were processed using FastQC (version 0.11.2.) to perform quality control on the raw sequences. The QIIME pipeline (version qiime2-2020.8) was used on 16S rRNA gene raw sequences from microbial communities. The pipeline includes: importing the files, demultiplexing using the q2-demux plugin, denoising, dereplication and removing chimeras using dada2 algorithm, de novo OTU clustering using vsearch with 97% identity, taxonomy assigning using classify-sklearn naïve base classifier against GreenGenes, Silva138 and DTDB databases, filtered to exclude mitochondria and chloroplast to avoid any possible contamination, phylogeny tree generating. Downstream analysis was performed using phyloseq (version 1.30.0), R/bioconductor package for handling and analysis of high-throughput phylogenetic sequence data. The taxonomy was first cleaned and filtered from empty taxas. The samples were then rarefied (using rarefy_even_depth function) at a minimum sequence depth and scaled to relative abundance.

The weighted-UniFrac distance matrix was calculated for samples in the experiment, i.e. alpha (estimate_richness function) and beta (distance function) diversity analysis (Bray-Curtis dissimilarities). For gut microbiome differential abundance, deseq2 (version 1.26.0), R/bioconductor package was used. This package allows differentially abundance analysis of the two groups before and after treatment. The filtered phyloseq object was converted into a DESeqDataSet object (using phyloseq_to_deseq2). PCA (principal component analysis) and heatmaps for samples distance were generated using Pheatmap (version 1.0.12). Rarified Scaled OTUs were labeled by lowest assigned taxa level possible and was summarized per taxa. Differential abundance was performed and significant taxa (|fc|>=1.5) selected at 0,3,6 weeks. To ensure that these significant taxa were not influenced by any potential contaminants, correlated taxa were compared with known lab contaminants mentioned by Salter and colleagues.

### Gut organ culture system

Fabrication of the gut organ culture device and gut organ culture experiments were performed as previously described (Yissachar et al., 2017). Briefly, intact whole colons were dissected sterilely from 14d old C57BL/6 mice reared under SPF conditions. The solid lumen content was gently flushed, and the gut fragment was threaded and fixed over the luminal input and output ports of the gut organ culture device, using a sterile surgical thread. The culture device was placed in a custom-made incubator that maintains temperature of 37°C, and tissue was maintained half-soaked in a constant flow of sterile medium using a syringe pump. Fecal samples of IBS patients, or purified bacterial cultures (*B. adolescentis* and *B. fragilis*) were resuspended in de-gassed tissue culture medium to a final concentration of 10^7^CFU/ml, and infused into the gut lumen using a syringe pump. Gas outlet in the device lid enabled flow of humidified and filtered, medical grade 95% O2/5% CO2 gas mixture into the device. Experiments were terminated at 2h post-colonization (for fecal samples) or 4h (for specific bacterial species), and tissue was subjected to further analysis.

### RNA sequencing and data analysis

Colon tissue fragments (3 mm) were stored in RNAlater overnight at 4°C, homogenized using a bead beater, and bulk RNA was extracted using RNeasy Plus Universal Mini QIAGEN kit. The Illumina TruSeq stranded mRNA library prep kit (#20020594) was used for library generation, and next generation sequencing was performed in the Bar-Ilan University scientific equipment center using the Illumina NextSeq platform (NextSeq 500 High Output v2 kit (#FC-404-2005)). For RNA sequencing data analysis, reads were aligned to the Mus musculus reference genome GRCm38 using STAR (version 020201), and quantification of reads was performed using htseq-count (version 0.12.4) and a list of genes (gtf file). Differential gene expression analysis was then performed using deseq2 (version 1.26.0), R/bioconductor package. DESeq2 applies the Wald’s test on estimated counts and uses a negative binomial generalized linear model to determine differentially expressed genes and the log-fold changes. PCA and heatmaps for samples distance were generated for data visualization using Pheatmap (version 1.0.12). Significant differentially expressed genes were selected using threshold values of *p*-value smaller than 0.05 and log2 fold change greater or equal to 1. The data was further transformed, and the normalized values were extracted using regularized logarithm (rlog) to remove the dependence of variance on mean. For pathway enrichment analysis, Metascape (metascape.org) was used to analyze differentially expressed genes. Additionally, Gene Set Enrichment Analysis (GSEA version 4.0.3) was used for all the genes ranked (-log10(pvalue) /sign(log2FoldChange)), converted to Human genes, using three datasets: hallmark, Curated Canonical Pathways and GO gene sets. GSEA results were categorized and plotted using ggplot2 (version 3.3.2). Next, gut microbial taxa (7 taxa) were correlated with colonic significant differentially expressed genes (206 genes) from their respective overlapping samples (8), using Pearson correlation on significant gene expression normalized counts and microbial taxa normalized abundance using corr.test function for each pair of gene-taxa. Significant gene-taxa correlations (*p*-value<=0.05) were visualized using corrplots (version 0.84), where the strength of the correlation is indicated by the color and size of the visualization element (square) and the significance of the correlation is indicated via asterisk.

### Antibiotic treatment and oral gavage

For broad range antibiotic treatment, 1g/L ampicillin sodium salt (Fisher Bioreagents), Metronidazole (Acros Organics), Neomycin sulfate (Fisher Bioreagents) and 0.5g/L Vancomycin hydrochloride (Acros Organics) were dissolved in normal drinking water. SPF mice received antibiotic treatment for 7d, and then were orally gavaged with 100µl of bacterial culture (*B. adolescentis* or *B. fragilis*; 10^9^CFU/ml) or sterile PBS daily, for 3d.

### Cell cultures

Human colon colorectal adenocarcinoma (CaCo-2) cell line was kindly provided by Prof. Ohad Gal Mor (Sheba Medical Center, Israel). The cells were cultured at 37°C and 5% CO2 until reaching approximately 80% confluence and were then sub-cultured using a trypsin-EDTA solution. The cells were maintained in Dulbecco’s modified Eagle medium (DMEM) and supplemented with 20% heat inactivated fetal bovine serum (FBS), 2 mM L-glutamine (L-Gln) and 100 U/ml Penicillin/Streptomycin. For co-culture experiments, CaCo-2 cells were cultured with growth media lacking Penicillin/Streptomycin and were seeded on a 24-well plate (2x10^5^ cells per well) or on 12-well plate (at 4x10^5^ cells per well). The cells were grown for 4d and then incubated for 4h at 37°C, 5% CO2 in the presence of bacterial cultures, resuspended in CaCo-2 growth media at 10^7^ to 10^8^ CFU/ml.

### RNA extraction and RT-PCR

RNA was extracted from CaCo-2 cells using Direct-zol™ RNA Microprep (ZYMO cat: R2062), according to manufacturer’s instructions. The concentration and absorbance at 260 nm and 280 nm were measured to assess RNA purity. RNA was reverse transcribed into cDNA using qSqript (Quantabio, cat:95047-100). SYBER green (ThermoFisher scientific cat: 4385614) was used to evaluate gene expression, using a real-time PCR apparatus (CFX 96, Bio-Rad) system. The primers used are listed in below, using eEF2 as a reference gene. Relative gene expression levels were determined using the 2^−ΔΔCT^ method.

The primers used for RT-PCR:

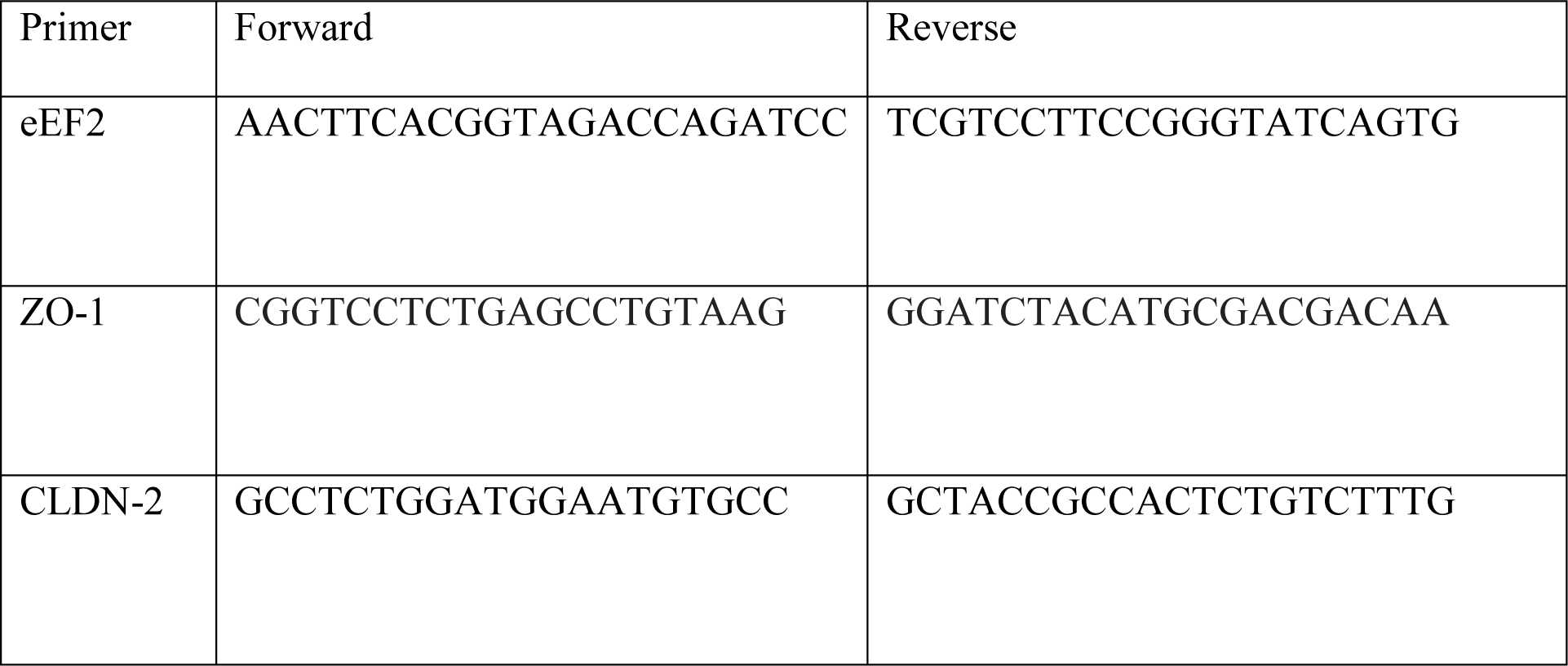

### Immunofluorescence staining and confocal microscopy

CaCo-2 cells growth on cover glass were fixed with cold methanol for 10 minutes. After washing with PBS, cells were incubated with blocking solution (10% goat serum, 0.1% BSA, 0.3% glycine 0.1% tween in PBS) for 1h. The cells were then labeled with primary antibody against ZO- 1 (BD cat:610967) overnight at 4°C. Excess antibody was washed 3 times with the blocking solution and the cells were incubated with secondary antibody (Cy5, Jackson cat: 715-175-151) for 1h at room temperature. Coverslips were washed and mounted using an antifade reagent (SIGMA F6182). Cells were visualized using a confocal fluorescence microscope (Leica) and processed and analyzed using ImageJ Software. For the tissue sections staining, intestinal tissues were embedded in OCT and stored at −80°C. The fresh-frozen tissues were sliced using a cryostat to 7 µm sections. The sections were fixed with cold acetone, washed twice in PBS and blocked (10% donkey serum, 0.1% triton in PBS) for 1h at room temperature. Tissues section were then stained for ZO-1 (Invitrogen cat: 40-2200), overnight at 4°C. Excess antibody was washed 3 times and incubated with secondary antibody (Cy3 anti rabbit) for 1h at room temperature. After washing the section were stained with DAPI and mounted using an antifade reagent (SIGMA F6182).

### Image analysis

In order to measure the ZO-1 intensity only in the periphery of the intestinal epithelium, an ImageJ (FIJI) macro was employed, using the following workflow: The images were first Gaussian smoothed (sigma=2), and then threshold and segmented by the Percentile auto threshold algorithm using the E-cadherin staining. Binary holes were filled to get a uniform layer. Next, an ROI (region of interest) band, representing the edge of the intestinal crypt, enclosing the ZO-1 staining, was created as follows: First, the segmented tissue was duplicated and eroded to get a thinner segment. Second, the segmented tissue image was duplicated and dilated to get a wider segment. Finally, the eroded image was subtracted from the dilated image to yield a band, representing the tissue edge. Note, that while the macro allows adjustment of the number of pixels to erode and dilate, once optimal numbers were determined for our images to cover ZO-1, these factors were kept constant for all treatments and for each experiment type. Finally, Analyze Particles was employed to measure the intensity of the total ZO-1 (Cy3) in the ROI.

The intensity of ZO-1 in cultured cells were measured as followed: Smoothing on the ZO- 1 image (channel 3 Cy3) using Gaussian blur (sigma=1) was applied. Next, the background was subtracted (rolling ball=10). The cells were then threshold using the automatic Otsu algorithm. Finally, the intensity of the Zo-1 marker was calculated according to the binary segmented image.

### Dynamics Trans-Epithelial Electrical resistance (TEER) measurements

CaCo-2 cells were seeded on a 96-well ‘impedance’ plate (5*10^5^ cells per well) (Axion BioSystems; cat: Z96-IMP-96B-25) and monitored by Maestro Edge platform (Axion BioSystems) until reaching full confluence. The barrier index (which represents barrier integrity) was calculated by Axion ‘Impedance’ module, as the ratio between cellular resistance at low frequency (1Hz) vs. high frequency (41Hz). The barrier index was measured at high temporal resolution of 1 minute and was normalized to the reference time (t=0). Additional correction was performed by normalizing to the barrier index of unstimulated cells.

### Scanning electron microscope (SEM)

Small intestinal tissue was fixed in 2.5% (wt/vol) paraformaldehyde (PFA), 2.5% glutaraldehyde, and PBS for 2 h at room temperature and then left overnight at 4°C. After fixation the samples were washed 3 times with PBS, samples were further processed by dehydration in graded levels of ethanol solutions of 50%, 70%, 95% and 100%, and then underwent dried with hexamethyldisilazane. The samples were then coated with gold and images were acquired on an Emission Scanning Electron Microscope (Quanta FEG 250, FEI).

## Acknowledgements

We thank Ron Unger for helpful advice, Sara Gerstein for help with organ cultures, the Mathis/Benoist lab for microbial strains, Shani Brown for critical editing of the manuscript and members of the Yissachar lab for insightful discussions. This work was supported by the Israel Science Foundation (grant No. 3114831), the Israel Science Foundation — Broad Institute Joint Program (grant No. 8165162), and the Gassner fund for medical research, Israel.

## Supplementary figures

**Supplementary Fig. 1.**
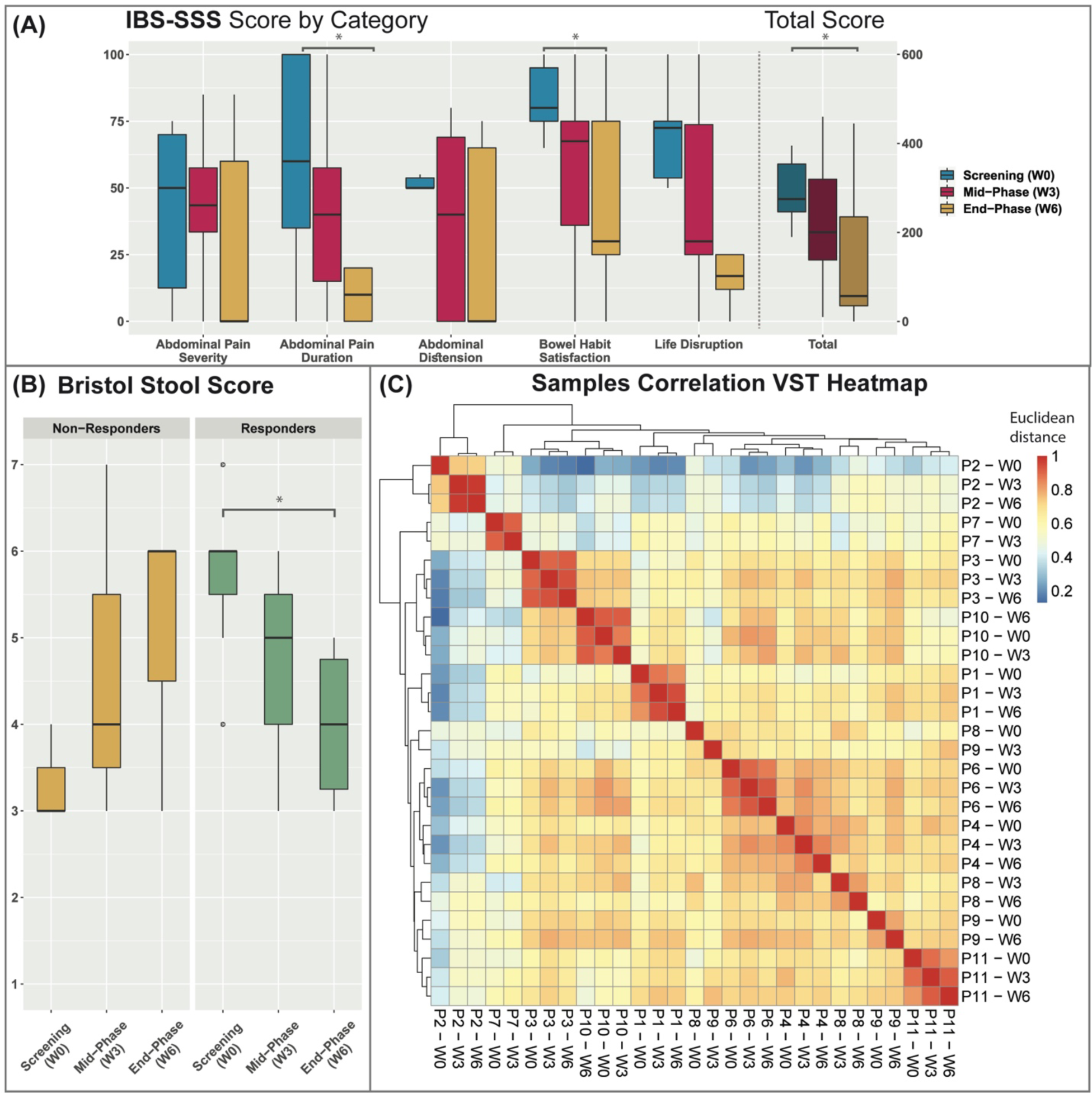
Clinical and microbial outcomes of low-FODMAP diet in IBS patients. **(A)** Changes in IBS symptom severity scale (IBS-SSS), summarizing survey scores filled by patients at each timepoint. Significant (* *p*<0.05) decrease (improvement) in scores was observed in two of the six survey categories comparing pre- (W0) and post-diet (W6): abdominal pain duration (37.9 ± 12.8) and bowel habit satisfaction (37.9 ± 13.8). Total score (sum of the six categories) showed significant improvement as well (146.3 ± 51.3; * *p*< 0.05). **(B)** Changes in Bristol Stool Score across timepoints divided into responders (improvement in IBS-SSS scores between starting and end point) and non-responders. Statistical significance was determined by ANOVA for repeated measures, with FDR post-HOC test. * *p* < 0.05) **(C)** Gut microbiome composition across all participants clustered first by patient, then by week on low-FODMAP diet. A heatmap of Euclidean distance matrix provides an overview of similarities and dissimilarities among samples. Each sample is OTUs abundance grouped data. The samples were transformed using Variance Stabilizing Transformation function, yielding a matrix of values that are approximately homoskedastic (having constant variance along the range of mean values).

**Supplementary Fig. 2.**
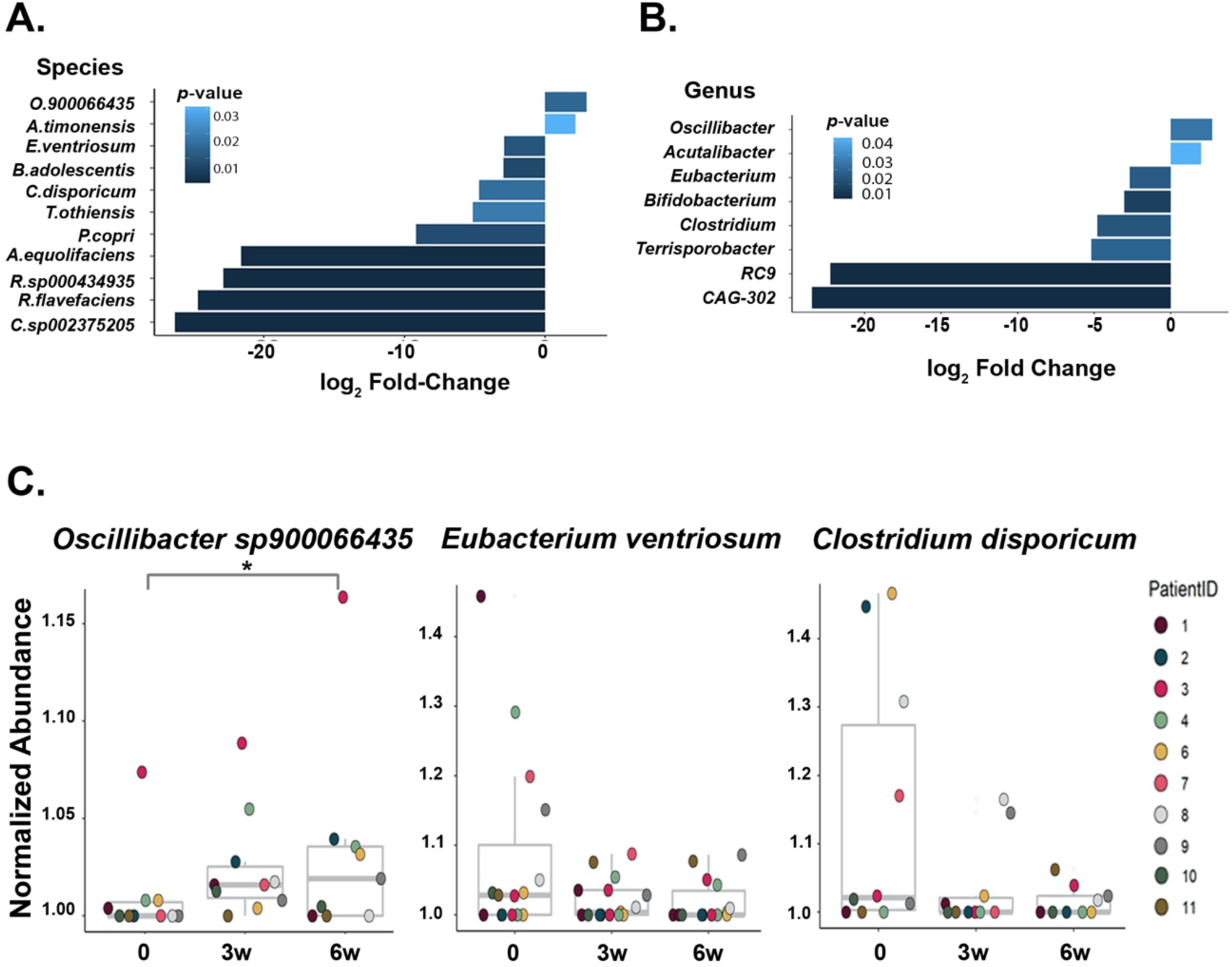
Differential abundance of gut microbiome following low-FODMAP diet. **(A-B)** Relative abundance of microbial taxa at the species **(A)** or genus **(B)** level (by DESeq2, Fold Change>1.5, Wald test *p*<0.05) in post-diet microbiota (6w), compared with pre-diet microbiota of IBS patients (n=10). **(C)** Normalized abundance of *Oscillibacter sp900066435, Eubacterium ventriosum* and *Clostridium disporicum* in patients’ gut microbiota along initial (0w), mid-phase (3w) and end-phase (6w) of low-FODMAP diet.

**Supplementary Fig. 3.**
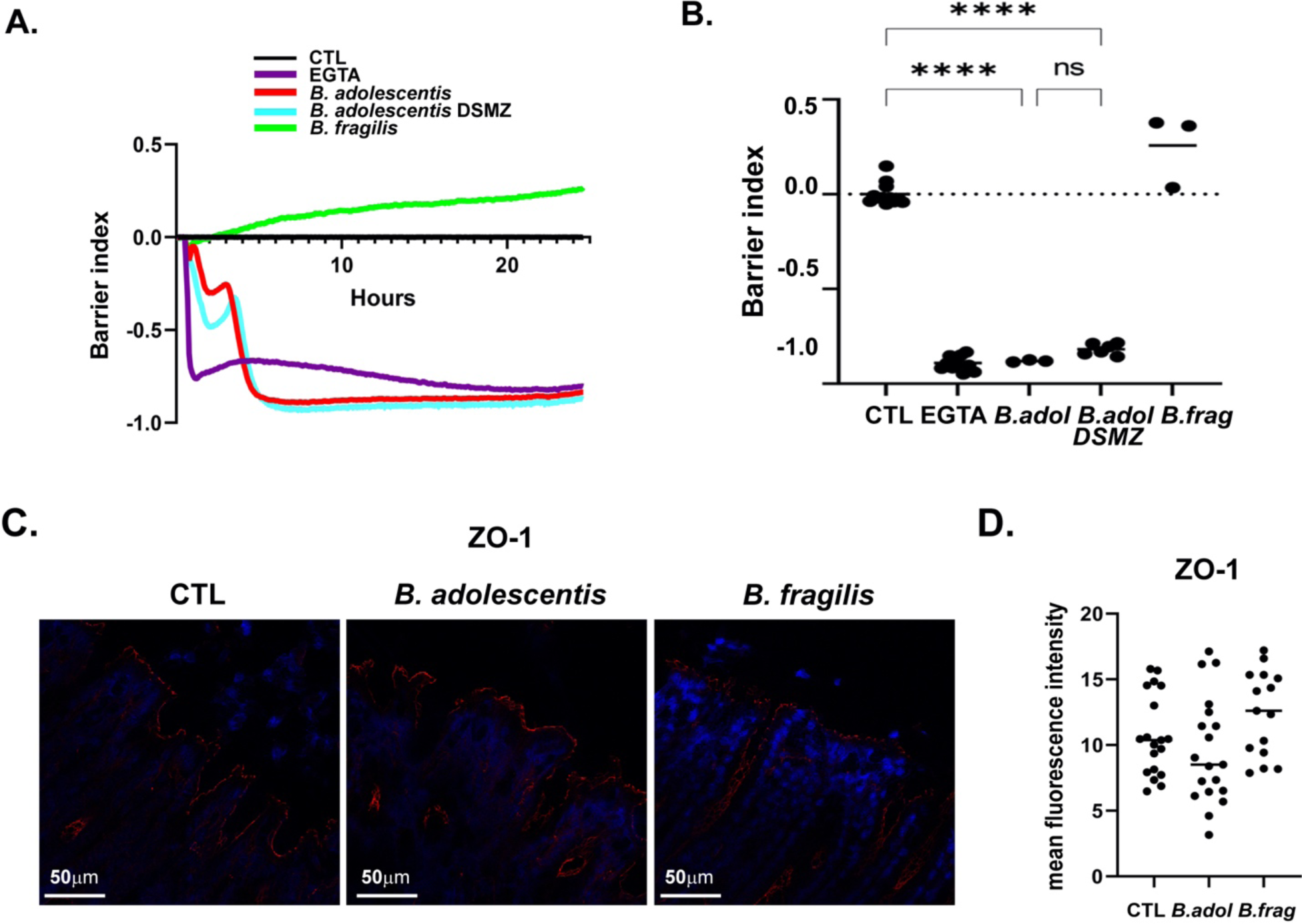
TEER assay using B. adolescentis acquired from different sources. **(A, B)** Normalized TEER values vs. time (**A**) and after 24h (**B)**. **(C,D)** Confocal microscopy images stained for ZO-1 (red, DAPI nuclear stain in blue) (**C**) and quantification of Mean Fluorescence Intensity (MFI) (**D**) of colon tissue from mice following oral gavage of *B.adolescentis* (B.adol), *B.fragilis* (B.frag) or sterile PBS (CTL).

